# Differential paralog divergence modulates evolutionary outcomes in yeast

**DOI:** 10.1101/063248

**Authors:** Monica R. Sanchez, Aaron W. Miller, Ivan Liachko, Anna B. Sunshine, Bryony Lynch, Mei Huang, Christopher G. DeSevo, Dave A. Pai, Cheryl M. Tucker, Margaret L. Hoang, Maitreya J. Dunham

**Author notes:** Current address: Zymergen, Emeryville, California, United States of America (Aaron W. Miller). Current address: Children's Hospital of Pittsburgh, Pittsburgh, Pennsylvania, United States of America (Bryony Lynch). Current address: Harborview Medical Center, Seattle, Washington, United States of America (Mei Huang). Current address: Exelixis, Dallas, Texas, United States of America (Christopher G. DeSevo). Current address: BioNano Genomics, San Diego, California, United States of America (Dave A. Pai). Current address: Wall High School, Wall Township, New Jersey, United States of 28 America (Cheryl M. Tucker). Current address: Nanostring, Seattle, Washington, United States of America (Margaret L. Hoang). Corresponding author (MD).

## Abstract

Evolutionary outcomes depend not only on the selective forces acting upon a species, but also on the genetic background. However, large timescales and uncertain historical selection pressures can make it difficult to discern such important background differences between species. Experimental evolution is one tool to compare evolutionary potential of known genotypes in a controlled environment. Here we utilized a highly reproducible evolutionary adaptation in *Saccharomyces cerevisiae* to investigate whether other yeast species would adopt similar evolutionary trajectories. We evolved populations of *S. cerevisiae, S. paradoxus, S. mikatae, S. uvarum*, and interspecific hybrids between *S. uvarum* and *S. cerevisiae* for ~200-500 generations in sulfate-limited continuous culture. Wild-type *S. cerevisiae* cultures invariably amplify the high affinity sulfate transporter gene, *SUL1*. However, while amplification of the *SUL1* locus was detected in *S. paradoxus* and *S. mikatae* populations, *S. uvarum* cultures instead selected for amplification of the paralog, *SUL2*. We measured the relative fitness of strains bearing deletions and amplifications of both *SUL* genes from different species, confirming that, converse to *S. cerevisiae, S. uvarum SUL2* contributes more to fitness in sulfate limitation than *S. uvarum SUL1*. By measuring the fitness and gene expression of chimeric promoter-ORF constructs, we were able to delineate the cause of this differential fitness effect primarily to the promoter of *S. uvarum SUL1*. Our data show evidence of differential sub-functionalization among the sulfur transporters across Saccharomyces species through recent changes in noncoding sequence. Furthermore, these results show a clear example of how such background differences due to paralog divergence can drive significant changes in evolutionary trajectories of eukaryotes.

## Author summary

Both comparative genomics and experimental evolution are powerful tools that can be used to make inferences about evolutionary processes. Together, these approaches provide the opportunity to observe evolutionary adaptation over millions of years where selective history is largely unknown, and over short timescales under controlled selective pressures in the laboratory. We have used comparative experimental evolution to observe the evolutionary fate of an adaptive mutation, and determined to what degree the outcome is conditional on the genetic background. We evolved several populations of different yeast species for over 200 generations in sulfate limited conditions to determine how the differences in genomic context can alter evolutionary routes when challenged with a nutrient limitation selection pressure. We find that the gene encoding a high affinity sulfur transporter in most species of the *Saccharomyces* clade becomes amplified, except in *S. uvarum*, in which the amplification of the paralogous sulfate transporter gene *SUL2* is recovered. We attribute this change in amplification preference to mutations in the non-coding region of *SUL1*, likely due to reduced expression of this gene in *S. uvarum*. We conclude that the evolutionary route taken by each organism depends on the genomic context, even when faced with the same environmental condition.

## Introduction

Understanding how organisms adapt to their environment is a fundamental goal of evolutionary biology. This goal has been complicated by the dependence on the reconstruction of historical events to make inferences about selective pressures and evolutionary mechanisms. Furthermore, it can be difficult to pinpoint genetic variation that causes new phenotypes of interest amid very divergent genomes. One approach to circumventing this limitation is to study evolution in the laboratory, where growth, environment, and population parameters can be controlled and dynamic adaptation events can be followed in real time [1–5]. However, experimental evolution has its own limitations, such as being too far removed from natural environmental factors and extending over only limited time scales. Merging laboratory evolution and comparative genomics could provide a more comprehensive view of processes that underlie evolution. In addition, comparative experimental evolution allows us to determine to what degree genetic background may result in differential functional innovation in the future [4,6].

One source of genetic novelty that may vary across divergent species is gene duplication. Gene duplicates can have different fates, either through dosage effects of an extra copy, splitting ancestral sub-functions or regulatory patterns over duplicates (sub-functionalization), or acquiring novel function (neo-functionalization) [7,8].

Alternatively, they can provide genetic redundancy to endow organisms with mutational robustness [9–11]. Duplications occur frequently during evolution and are commonly linked to genome innovations that result in an adaptive or phenotypic change to a particular environment [12,13]. After a duplication event, adaptation may result through the accumulation of mutations in the non-coding or protein coding regions of the genome, which may alter gene function, protein-protein interactions, or expression profiles. Accumulation of mutations in the coding region of each paralog may potentially modify active sites, affecting biochemical functionality, or alter binding interfaces and thus their interaction specificity [14]. Mutations in the non-coding region of each paralog may cause regulatory interactions in networks to be lost or re-wired, potentially leading to expression divergence between paralogs [15–17].

The *Saccharomyces sensu stricto* clade of species provides a particularly appealing platform for comparative studies of gene function. The last common ancestor of this group existed approximately 20 million years ago, with approximately 80% identity in coding sequences between *S. cerevisiae* and *S. uvarum* [18]. The *Saccharomyces* species are experimentally tractable, have high quality genome sequences [19,20], contain largely syntenic chromosomes [21], and can mate to form hybrids, including with the laboratory workhorse *S. cerevisiae*, providing access to a huge knowledge base and extensive toolkit of genetic and genomic resources. Additionally, the *Saccharomyces* genus is a result of a well-studied WGD event, which occurred just before the separation of *Vanderwaltozyma polyspora* from the *S. cerevisiae* lineage [22] and was itself probably a result of a hybridization event [23].

In this study, we compared the evolutionary outcomes upon sulfate-limited growth in chemostat culture between *S. cerevisiae, S. paradoxus, S. mikatae, S. uvarum*, and *S. cerevisiae/S. uvarum* hybrid strains and used whole genome sequencing and species-specific microarrays to identify resultant genetic changes. We discovered differential amplification of sulfate transporter gene paralogs *SUL1* and *SUL2* in the different species. The species-specific amplification preference correlated with the selective effects of amplification and deletion of each sulfate transporter gene. Analysis of functional divergence of the two paralogs across these species provides evidence for differential sub-functionalization between the *SUL1* and *SUL2*paralogs of *S. cerevisiae* and *S. uvarum*, driven largely by lineage-specific acquired changes in the non-coding region of *SUL1* in *S. uvarum*. In this work, we discovered an example of recent paralog divergence between two gene duplicates with altered gene expression between *S. cerevisiae* and *S. uvarum*, and demonstrated that such differences can alter the mechanisms by which these species adapt to future challenges.

## Results

### Adaptation through differential gene amplification: *S. cerevisiae* and *S. uvarum* species amplify different sulfate transporter genes

As described previously [24–26], evolved clones of *S. cerevisiae* selected during longterm continuous culture under sulfate-limitation reproducibly carry amplification events on chromosome II containing the high affinity sulfur transporter gene *SUL1* (representative event shown in **Fig 1A**). This mutation confers one of the highest (20-40% increase) and most reproducible (18/18 populations) fitness advantages known in the experimental evolution literature [24–26]. In order to determine whether other yeast species would follow this same evolutionary path, we performed two evolution experiments with a sister species, *S. uvarum*, in chemostats using the same condition in which the *SUL1* amplification has been observed for *S. cerevisiae*. Each experiment was initiated with a prototrophic diploid *S. uvarum* strain that had never before been exposed to long-term sulfate limitation in the laboratory (see materials and methods).

**Fig 1.**
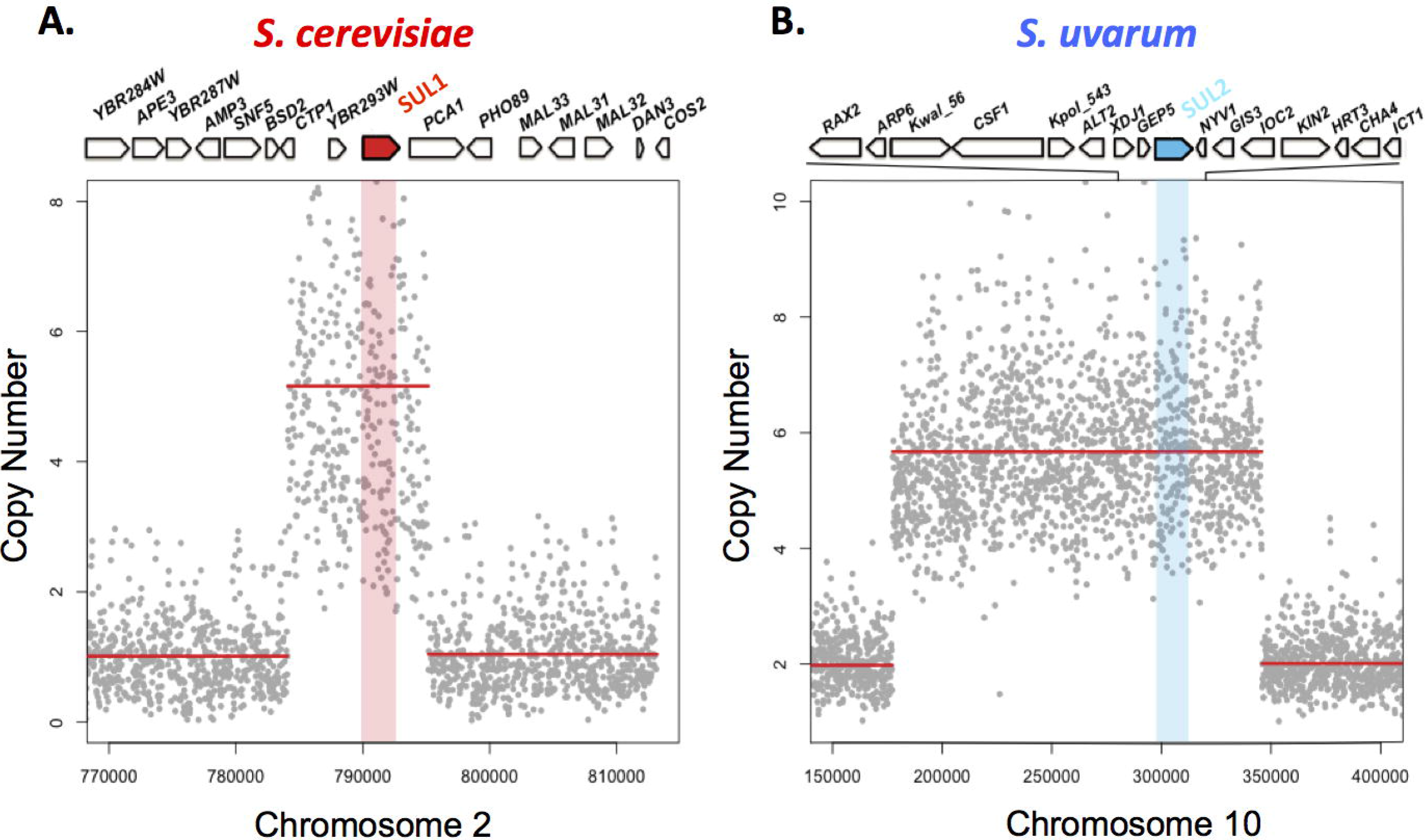
Fig 1. Adaptation through differential gene amplification between *S. cerevisiae* and *S. uvarum*. A) Copy number of relevant genomic segment in a representative evolved strain of *S. cerevisiae*. Copy number plots were calculated by sequencing-depth ratios between evolved and parental genomes in *S. cerevisiae* at 188 generations. Gray dots represent the per nucleotide read-depth averaged across 25bp windows. Segmentation-derived regions of equal copy number are indicated in red. Segmentation defines an ~11kb region with a copy number of 5. The region of the sulfate transporter gene *SUL1* gene is shaded red. B) Sequencing-depth ratios between evolved and parental genomes in an *S. uvarum* clone isolated at 510 generations. A large segmental amplification of an evolved clone defining a ~168kb region with a copy number of 5 includes the locus containing the sulfate transporter gene *SUL2*, shaded in blue. Genes aligned at the top represent the loci in the expanded panel.

In contrast to the amplification of *SUL1* in the *S. cerevisiae* clones, no amplification of this locus was observed in the two populations of *S. uvarum* evolved under sulfate limitation for 500 generations. However, the locus containing the gene *SUL2* was amplified in both populations as determined through microarray-based comparative genomic hybridization (aCGH) (**S1 Fig**). Two clones from one population were analyzed further by deep sequencing, revealing an internal segment of chromosome X containing the gene *SUL2* at an increased copy number of 5 in one of the two clones (**Fig 1B**). The fitness benefit of this evolved clone was 20% when competed against the ancestral strain (n=4, (**Table 1**). Although the exact function of the protein Sul2 has never been experimentally tested in *S. uvarum*, Sul2 has been identified as a lower affinity transporter of sulfate in *S. cerevisiae* [27].

**Table 1.** Fitness coefficient of evolved clones.

| Species | Generations | Clones | Fitness Coefficient, % | SUL1 Copy Number |
| --- | --- | --- | --- | --- |
| *S. cerevisiae* | 210 | Clone1 | 44.2 ± 8.2 (n = 2) | 5 |
| *S. uvarum* | 510 | Clone1 | 21.8 ± 2.37 (n = 4) | 5 |

We next set out to explain the differential amplification of *SUL1* and *SUL2* in these closely related species. We hypothesized that the different evolutionary outcomes could result from divergence in gene function—Sul2 may be the higher affinity transporter gene in *S. uvarum* and so its amplification causes a higher fitness benefit—or from changes in chromosomal context that affect amplification rate or amplicon fitness. We test these hypotheses below.

### The genomic context of *SUL1* in *S. uvarum*

We hypothesized that the preference for the amplification of *SUL1* in *S. cerevisiae* could be due to changes in chromosomal context between the two species that might affect the propensity of the region to amplify. *SUL1* in *S. cerevisiae* is located on the right arm of chromosome II, near the telomere. The *S. uvarum* ortholog is located in a syntenic region. Adjacent sequences to the telomeric repeats, including X and Y’ elements as well as subtelomeric gene families, have been shown to be rapidly evolving across species of the *Saccharomyces* clade, possibly contributing to a difference in mutation rate [20,28–30]. Also, as compared with the *S. cerevisiae* genome, the left portion of this chromosome contains a reciprocal translocation with a region syntenic to the right arm of *S. cerevisiae* chromosome IV [21]. The regions immediately adjacent to *SUL1* are largely syntenic, though the gene just distal to *SUL1* in *S. cerevisiae, PCA1*, is missing in *S. uvarum*. In *S. cerevisiae*, this region also contains a DNA replication origin (ARS228), which we previously demonstrated to be involved in the generation of the amplification [31,32]. To test for replication origin function, we cloned the corresponding region from *S. uvarum* and tested it for the ability to support plasmid replication (i.e., an assay for Autonomously Replicating Sequences, or ARSs). Like *S. cerevisiae, S. uvarum* does contain an active ARS in this region. Furthermore, both ARSs appear to function as chromosomal replication origins based on density transfer experiments (Liachko *et al*. in preparation).

### *SUL1* in *S. uvarum* can be amplified 1 in the absence of *SUL2*

Although the origin of replication is present, there may be other differences near *SUL1* in *S. uvarum* that might explain why this region has not been observed to amplify in the evolved strains. To test if *SUL1* is capable of amplification, we evolved four haploid *sul2*Δ strains of *S. uvarum* in sulfate-limited media and tested the evolved populations for copy number variation using aCGH. At 260 generations, we identified an amplification of the *SUL1* locus in one of the four populations and no other amplifications in the other three populations (**Fig 2**). This result indicates that the *SUL1* locus in *S. uvarum* has the capacity for amplification, but does not attain high frequency in populations initiated with strains containing both *SUL1* and *SUL2* genes.

Alternatively, the *SUL2* locus cannot amplify in *S. cerevisiae*. To test if the *SUL2* locus can amplify in *S. cerevisiae*, we evolved four haploid strains of *S. cerevisiae* in which *SUL1* has been deleted (*sul1*Δ) in sulfate-limited media and tested the evolved populations for copy number variation using aCGH. We identified an amplification of the *SUL2* locus in all four populations indicating that *SUL2* can amplify in *S. cerevisiae*, but these amplifications do not attain high frequency in evolution experiments performed with strains in which the *SUL1* gene is present (S2 Fig). We note that these experiments leave open the possibility that differences in amplification rate might contribute to the observed differences in amplification propensity. We have so far been unable to measure the amplification rate of these loci and so have not tested this hypothesis. However, another possible explanation for these results is that *SUL2* amplification may have a greater selective effect in *S. uvarum*. To test this, we performed additional experiments to determine the functional contribution of each gene from both species.

**Fig 2.**
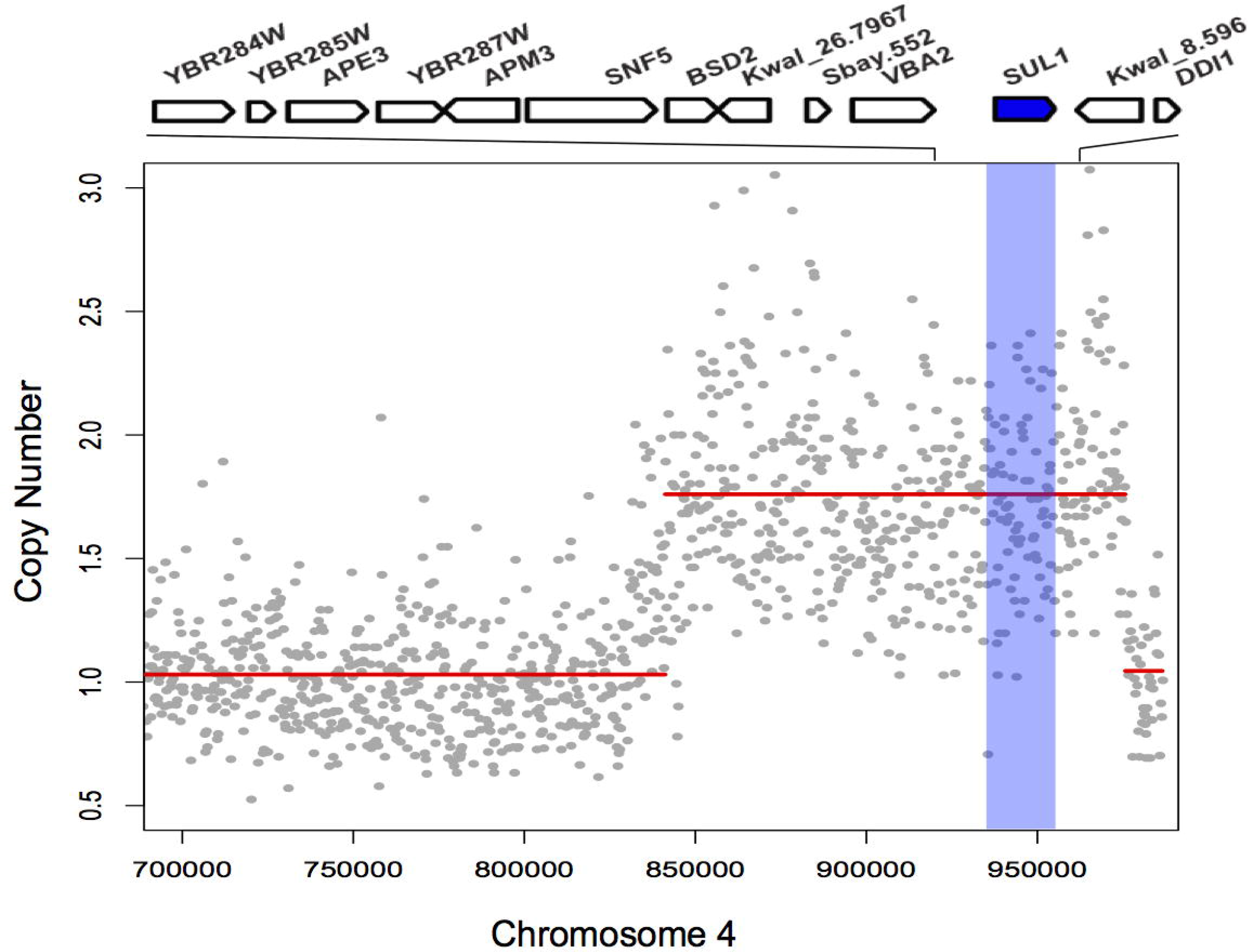
*SUL1* in *S. uvarum* can amplify but such amplifications are not observed when *SUL2* is expressed. Array CGH of evolved clone versus the parental genomes of a *sul2Δ S. uvarum* strain. Array data (gray dots) indicate a copy number amplification. Segmentation-derived regions of average copy number are indicated in red. Segmentation defines a ~134kb region of chromosome IV with a copy number estimation of 2. Genes along the top are represented in the locus in the expanded panel.

### Extra copies of sulfur transporter genes from *S. cerevisiae* and *S. uvarum* confer differential fitness effects

From comparisons with the reconstructed ancestral genome [33], *SUL2* appears to be the ancestral copy of the sulfur transporter, with *SUL1* being a more recent gene duplicate after a small-scale duplication (SSD) event. Amino acid conservation between *SUL1* and *SUL2* in *S. cerevisiae* is 62.5% and 61.3% shared identity in *S. uvarum*, whereas *SUL1* from *S. cerevisiae* and *SUL1* from *S. uvarum* share 84% identity and *SUL2* from *S. cerevisiae* and *SUL2* from *S. uvarum* share 87% identity, indicating that the sulfate transporter genes are correctly annotated.

To test whether the functions of these genes may have diverged between these species, we measured the fitness effects of having additional copies of each gene. Previous studies have shown that the addition of *SUL1* on a low copy plasmid in *S. cerevisiae* increases the fitness of the strains by ~40% [26]. To determine the effect of additional copies of *SUL1* and *SUL2* from *S. cerevisiae* and *S. uvarum*, we transformed *S. cerevisiae* with ARS/CEN plasmids individually containing each SUL gene along with 500bp upstream of the coding region. Fitness assays were also attempted in the *S. uvarum* background; however, the *S. cerevisiae* CEN plasmid we used was lost at a high rate, as previously observed [19], precluding clear results. We performed chemostat competition experiments between strains harboring additional copies of each gene in *S. cerevisiae*. The pairwise competitions provided fitness data that allowed us to more precisely determine the rank order of the fitness benefit of each gene amplification. The strain with an extra copy of *SUL1* from *S. cerevisiae (ScSUL1)* outcompeted all other strains, followed by *SUL2* from *S. cerevisiae (ScSUL2)*, which had a comparable fitness effect to *SuSUL2*. The strain with the *SuSUL1* gene had the lowest fitness effect of all genes tested (Fig 3). This result suggests that *SUL2* may have maintained a similar function between the two species, but *SUL1* function may have diverged. In support of our original hypothesis, the *SUL2* gene from *S. uvarum (SuSUL2)* conferred a greater fitness effect than the *S. uvarum SUL1 (SuSUL1)*. This is also consistent with our predictions based on the evolution experiments, suggesting that *SuSUL2* amplification may have a greater selective benefit than amplification of *SuSUL1*.

**Fig 3.**
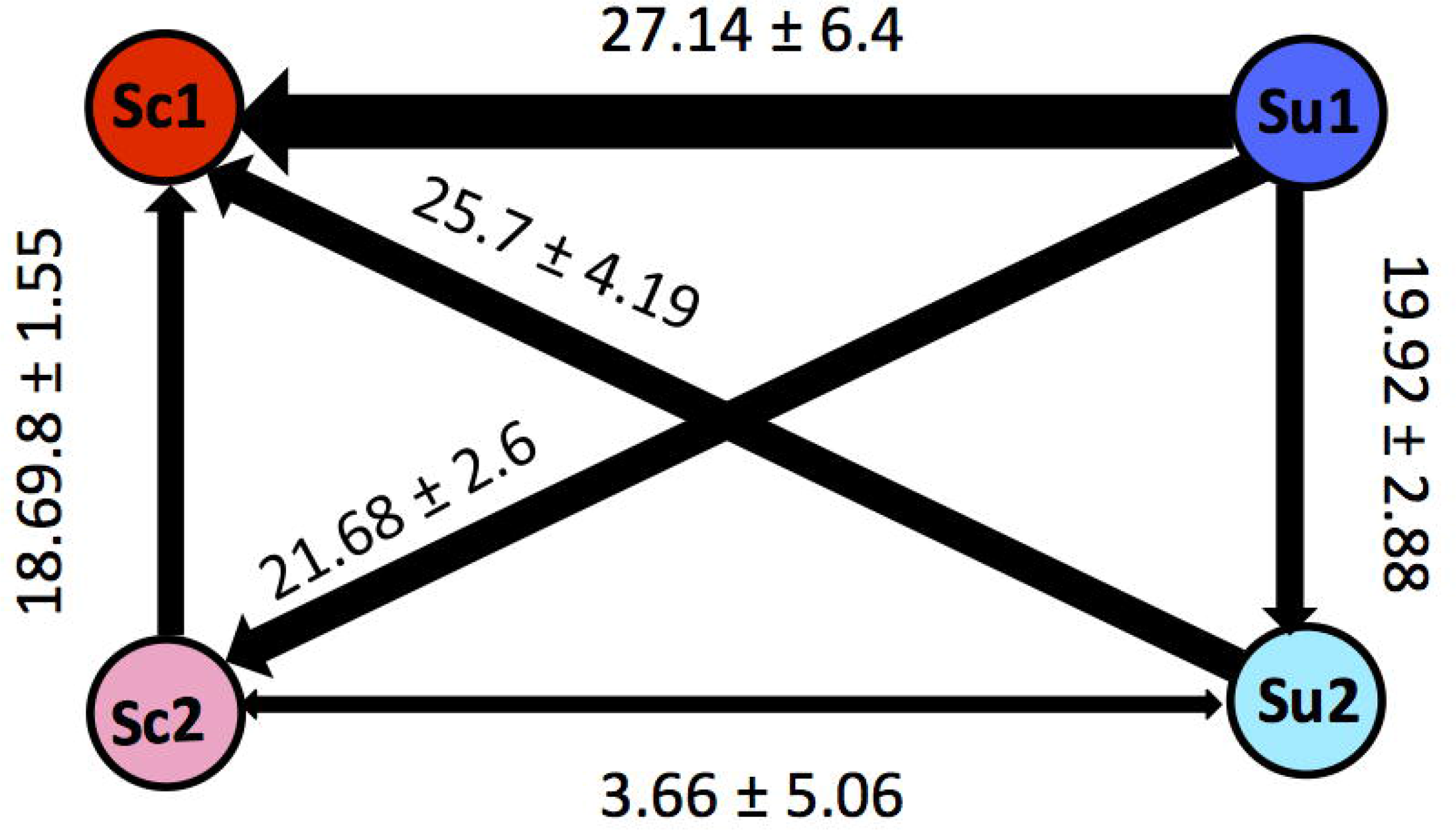
*SUL1* and *SUL2* have differential fitness effects between *S. cerevisiae* and *S. uvarum*. The relative fitness of four strains of *S. cerevisiae* containing CEN plasmids with a *SUL1* or *SUL2* allele from *S. cerevisiae* or *S. uvarum* was determined in a pairwise manner. The fitness was measured in the chemostat under sulfate-limited conditions against a GFP-marked lab strain also containing CEN plasmids with either *SUL1* or *SUL2* from *S. cerevisiae* or *S. uvarum*. Each arrow corresponds to the mean of 4 or more replicates ±SE. The thickness of the arrow illustrates the degree of fitness and the direction of the arrow points to the allele that outcompeted the other strain.

### *S. cerevisiae* × *S. uvarum* hybrid strains amplify *ScSULl* allele

In addition to testing the fitness effects of each *SUL1* and *SUL2* gene independently, we also investigated the amplification preference in the context of having all alleles present in one genome. Given the results from the single gene plasmid experiments above, we predicted that *ScSUL1* would be the preferred allele for amplification. We created *de novo S. cerevisiae/S. uvarum* hybrid strains and subjected them to hundreds of generations of growth in sulfate-limited continuous culture. Evolved strains were then analyzed by aCGH to determine differences in genome content from their ancestral strains.

Amplification of segments containing the *SuSUL1* or *SuSUL2* gene was never observed in 16 clones from 8 independent populations, and *SuSUL1* was even found deleted in one evolved clone, displaying loss of heterozygosity at this locus (**S3 Fig**). In contrast, the *S. cerevisiae* copy of *SUL1* was found amplified in 14/16 evolved clones (**Fig 4**). Copy numbers estimated from the array CGH data ranged from 3 to as many as 20 copies of *SUL1*. Centromere-proximal breakpoints varied from population to population, but amplicons extended to the most distal telomeric probe in all cases. Additional rearrangements were rarely observed in these strains (**S3 and S4 Figs**). When all four alleles are present in the same genome, *ScSUL1* amplifications are preferentially recovered, suggesting that *ScSUL1* amplification yields the greatest fitness advantage in this particular environment and genomic context.

**Fig 4.**
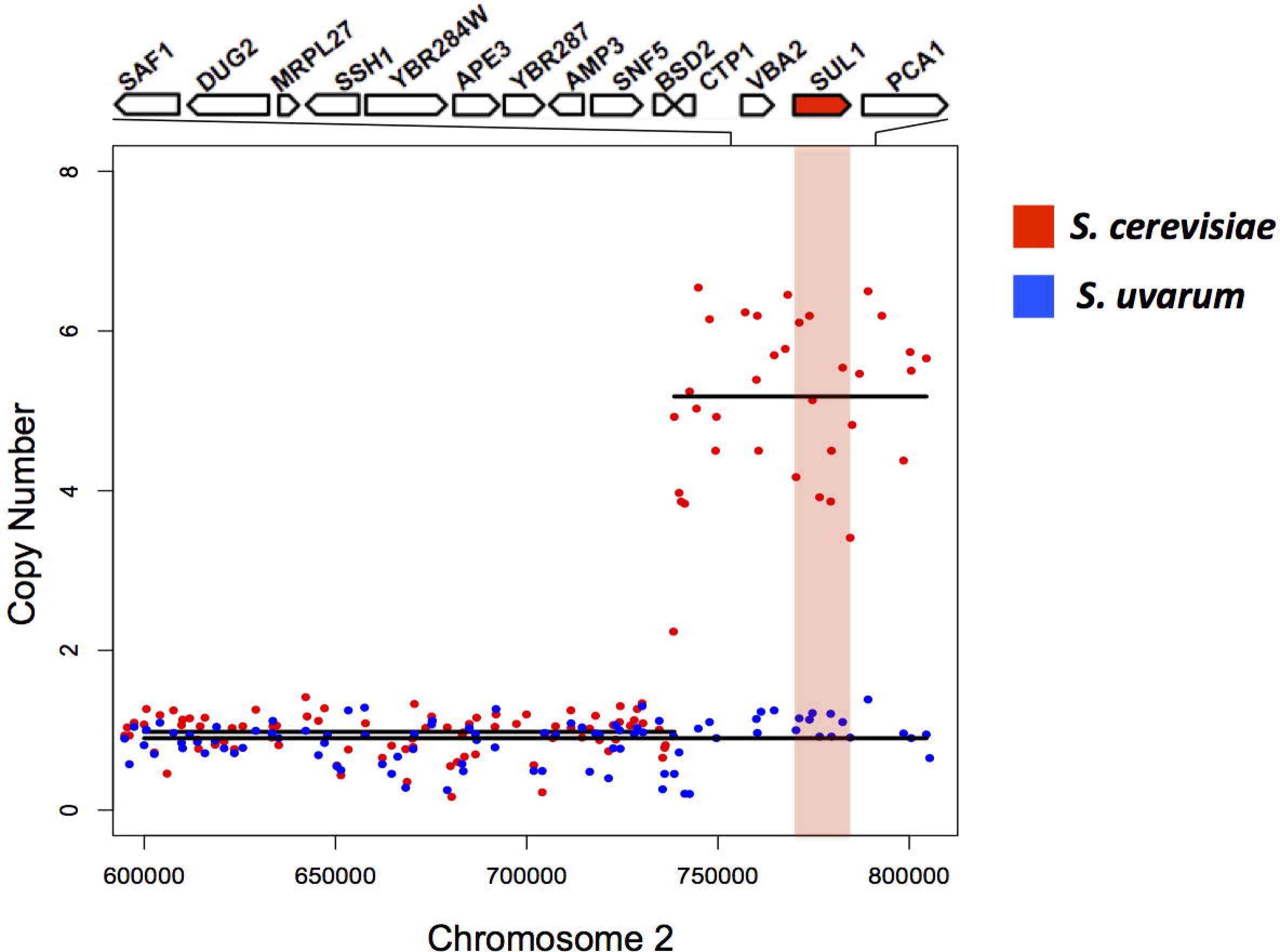
*S. cerevisiae* x *S. uvarum* hybrid strains amplify *S. cerevisiae SUL1*. Array CGH of evolved hybrid clone versus the parental *S. cerevisiae/S. uvarum* hybrid genome. Array data from *S. cerevisiae* (red dots) and *S. uvarum* /blue dots) support a copy number amplification of the *SUL1* locus of the *S. cerevisiae* allele. Black lines indicate the segmentation-derived regions of average copy number. Segmentation defines a ~65kb region of chromosome two with a copy number estimation of 5. The region of the sulfate transporter gene *SUL1* gene is shaded red.

### Deletion of sulfate transporter genes display differential effects between *S. cerevisiae* and *S. uvarum*

We have shown that the addition of extra copies of each gene results in an increased fitness in *S. cerevisiae*, with *ScSUL1* yielding the greatest fitness increase, a result that corresponds to the amplification preferences in evolved strains derived from an interspecific hybrid. Unfortunately, we were not able to test the fitness effect of plasmids carrying additional copies of *SUL1* vs. *SUL2* directly in *S. uvarum*. However, we were able to delete *SUL1* and *SUL2* in both *S. cerevisiae* and *S. uvarum* backgrounds to determine the relative fitness contributions of these loci in each background. We created *sul1Δ* and *sul2Δ* haploid strains and measured the competitive fitness of each null mutant in sulfate-limited conditions. We competed the *sul1*Δ and *sul2*Δ strains within each species against each other to calculate the fitness effect of each mutant. In *S. cerevisiae*, the *sul2*Δ strain outcompeted the *sul1*Δ strain, suggesting that *SUL1* in *S. cerevisiae* is the gene that is more important for growth in sulfate-limited conditions. Conversely, in *S. uvarum*, the *sul1*Δ strain out competed the *sul2*Δ strain, suggesting that *SuSUL2*, rather, is the gene that is more important for growth in sulfate-limited conditions (**Fig 5**). Taken together with the fitness data from increasing the copy number of each gene, these data suggest differential *SUL1* and *SUL2* fitness contributions across these two species despite the genes' similarity in amino acid composition.

**Fig 5.**
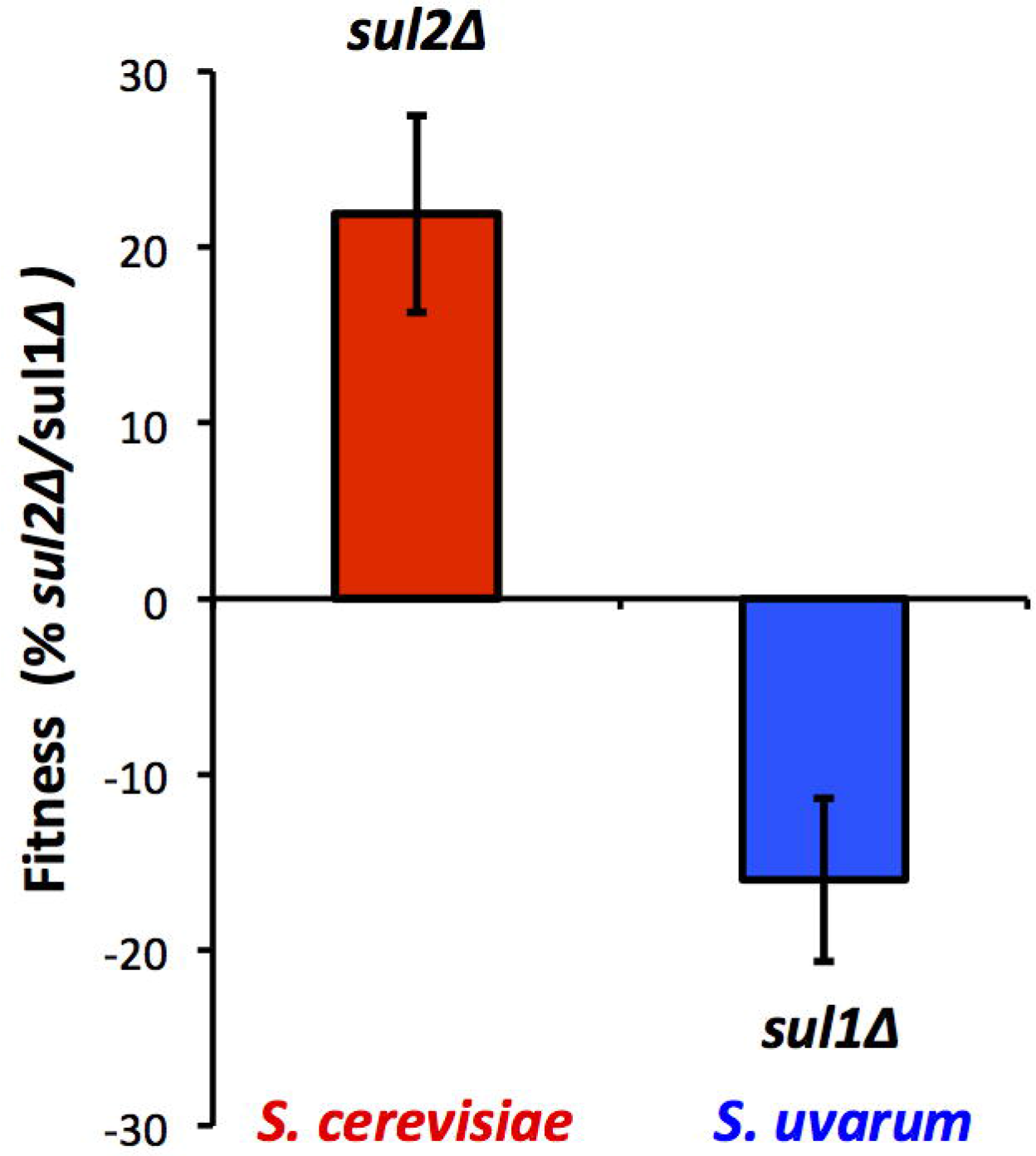
*sul1Δ* and *sul2Δ* have differential fitness effects between *S. cerevisiae* and *S. uvarum*. The relative fitness of two strains containing either a *sul1Δ* or *sul2Δ* allele in *S. cerevisiae* (red) and *S. uvarum* (blue) respectively was determined in a pairwise manner within each species. The fitness was measured in the chemostat under sulfate-limited conditions against a strain containing *sul2Δ* or *sul1Δ* allele in *S. cerevisiae* (red) and *S. uvarum* (blue) respectively. The proportion of each strain was determined by monitoring canavanine resistance, which differentially marked the competing strains. Each bar corresponds to the mean of 6 or more replicates ±SD. In *S. cerevisiae, sul2Δ* outcompetes *sulΔ* and in *S. uvarum sulΔ* outcompetes *sul2Δ*.

### *SUL1* amplification in other species of the *sensu stricto* clade

In order to determine where the divergence in relative fitness effects between *SUL1* and *SUL2* in *S. cerevisiae* and *S. uvarum* occurred in evolutionary history, we tested the fitness of *SUL1* and *SUL2* from *S. paradoxus* and *S. mikatae*—two other species of the *sensu stricto* clade—and *SUL2*from *Naumovozyma castellii*, a more distant species that has not undergone gene duplication of this locus. We cloned the genes along with 500bp upstream of the coding region from each species into an ARS/CEN plasmid and determined the relative fitness effect of the addition of the *SUL* genes in *S. cerevisiae* when competed against a plasmid-free strain. This experiment allowed us to calculate the relative fitness coefficient of each strain. All strains showed significantly higher fitness than wild type *S. cerevisiae*, with the relative fitness coefficients ranging from 12.7% to 37.2%. The *S. cerevisiae SUL1 (ScSUL1)* plasmid conferred the greatest fitness benefit of 37.2% (**Fig 6**). The strains containing *SUL1* from *S. paradoxus* and *S. mikatae* conferred a greater fitness advantage than *SUL2* from the respective species. In *N. castellii*, the singleton *SUL2* conferred a fitness advantage of 36.3% (**Fig 6**). These results suggest that the last common ancestor of *S. cerevisiae, S. paradoxus*, and *S. mikatae* may have acquired adaptive mutations in *SUL1*. Alternatively, the *S. uvarum SUL1* paralog may have acquired mutations that decreased its fitness only in that lineage.

**Fig 6.**
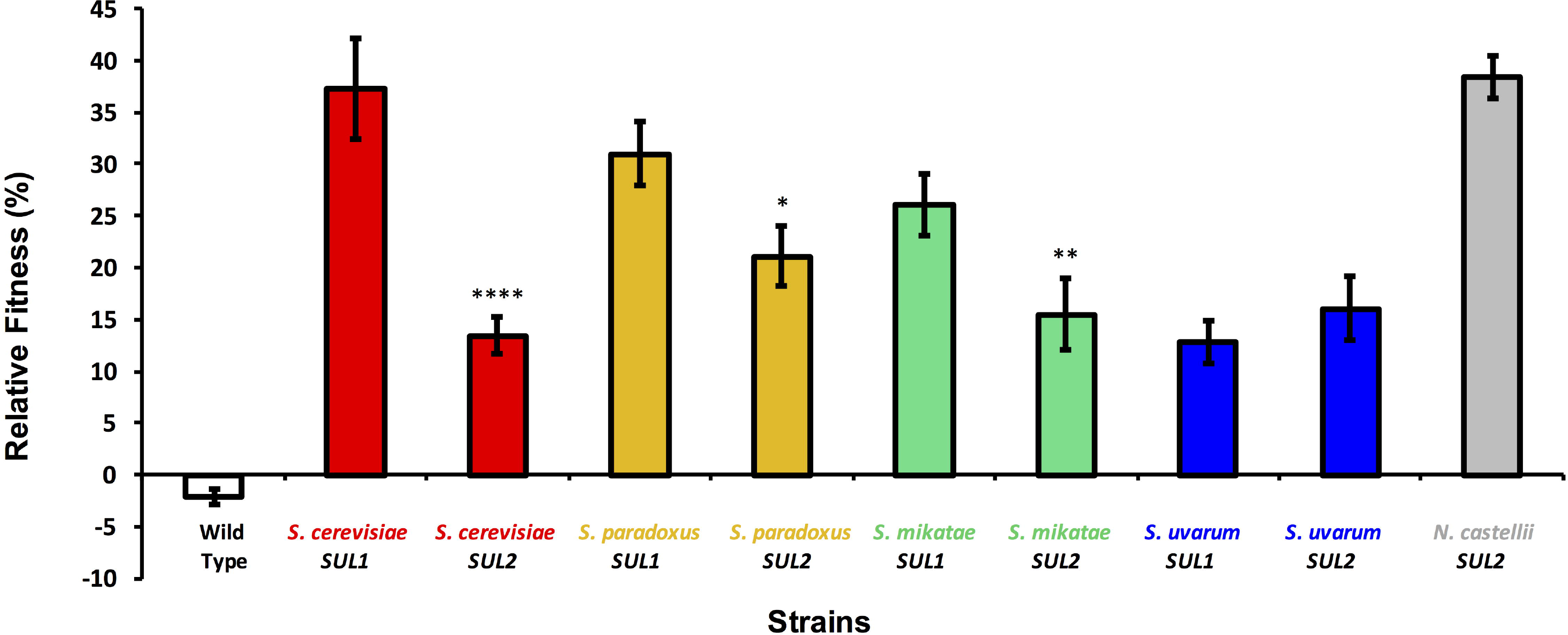
Plasmid-borne copies of *SUL1* cause a higher fitness benefit than *SUL2* in all species except *S. uvarum*. The relative fitness of 9 *S. cerevisiae* strains containing a plasmid with *SUL1* and *SUL2* alleles from *S. cerevisiae, S. paradoxus, S. mikatae, S. uvarum*, and *N castellii SUL2*. The fitness was measured in the chemostat under sulfate-limited conditions against a GFP-marked lab strain. Each bar corresponds to the mean of 4 or more replicates ±SD.

From these data, we can make predictions about the types of genomic events that would occur if we evolved *S. paradoxus* and *S. mikatae* under sulfate limited conditions. Since *SUL1* from both species resulted in the highest fitness benefit, we would expect to select for amplifications of the *SUL1* locus. To test this, we grew four populations of *S. paradoxus* and *S. mikatae* for 200 generations in sulfate limited chemostats and determined the copy number variation between evolved populations and each ancestral strain using deep sequencing. Consistent with expectations, we identified two populations with an amplification containing the *SUL1* locus in *S. paradoxus* and one population in *S. mikatae* (**Fig 7**). Other aneuploidy and segmental amplifications occurred in addition to the *SUL1* locus amplification in the evolved populations (**S5 and S6 Figs**); however, none of these copy number variants included the *SUL2* locus. Overall, these data are consistent with the previous gene function measurements of each allele in *S. cerevisiae*, indicating that *SUL1* is more adaptive when amplified in *S. paradoxus* and *S. mikatae*.

**Fig 7.**
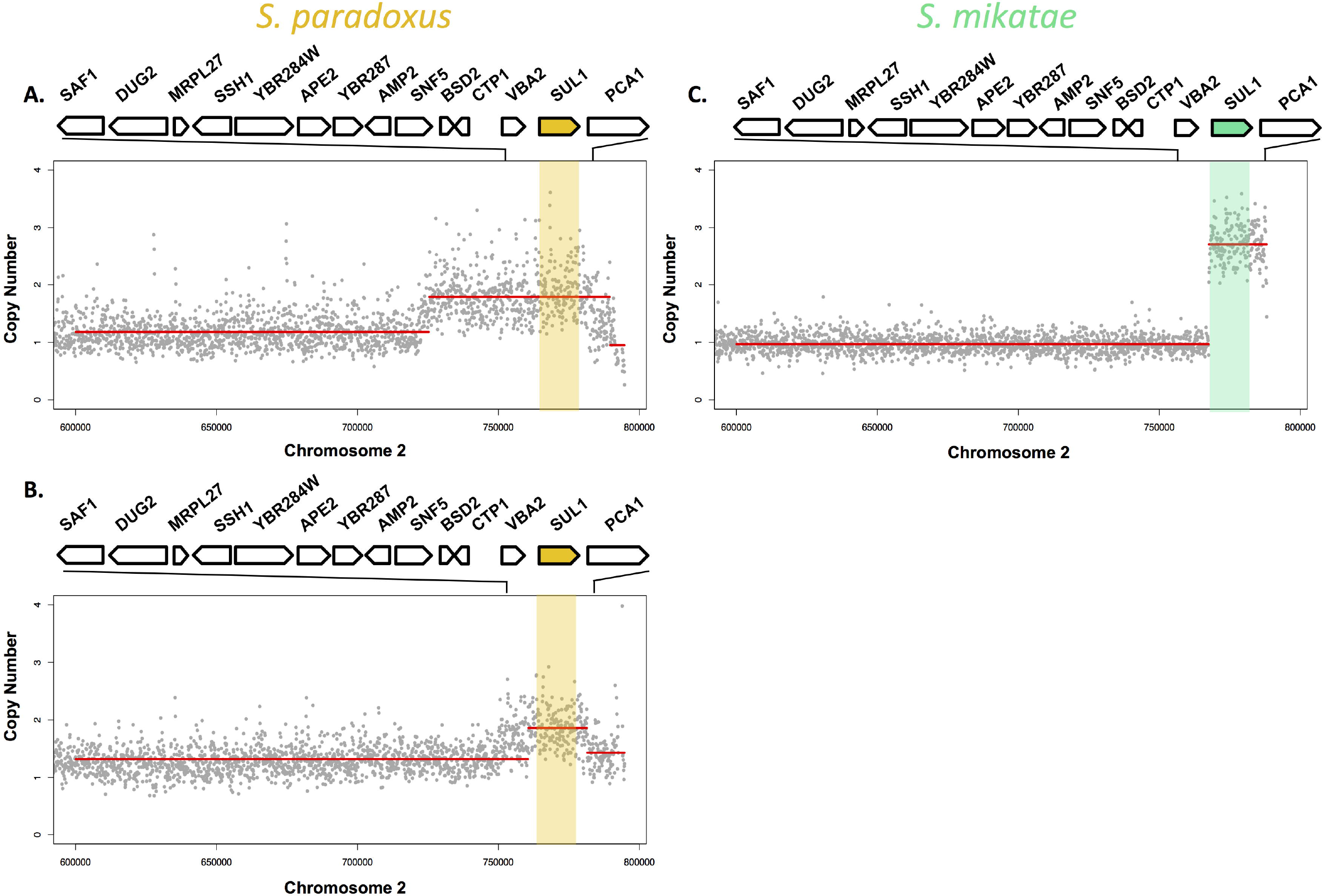
*SUL1* amplification in *S. paradoxus* and *S. mikatae* evolved populations. Copy number plots calculated with sequencing-depth ratios between evolved and parental genomes in 2 populations of *S. paradoxus* and one population of *S. mikatae* at ~200 generations. Gray dots represent the per nucleotide read-depth averaged across 100bp window and normalized against the average coverage of the ancestral strain. Segmentation-derived regions of average copy number are indicated as red lines. Genes aligned the top are represented in the locus in the expanded panel. The *SUL1* gene is shaded yellow in *S. paradoxus* and green in *S. mikatae*. A) Segmental amplification defines a ~64kb region with a copy number of 2. B) Segmental amplification defines a ~20kb region with a copy number of 2. C) Segmental amplification defines a ~20kb region with a copy number of 3.

### The species-specific relative fitness contributions among *SUL* genes are largely driven by promoter sequences

Based on the similar results across *S. cerevisiae, S. mikatae*, and *S. paradoxus*, we decided to focus on understanding what is different about the paralogs in *S. cerevisiae* vs. *S. uvarum*. To identify the genetic region responsible for the differences in fitness effects of *SUL1* and *SUL2* between the two sister species, we created chimeric constructs composed of different combinations of the promoter and open reading frame (ORF) of each gene. Rich *et al* recently used a deep mutational scanning approach to identify the functional elements of the *ScSUL1* promoter that are crucial for growth in sulfate limitation [34]. Based on their results, we cloned 500bp upstream of each ORF (the region encompassing all elements that positively influence *SULVs* fitness contribution) and cloned the ORF until the stop codon. We then cloned all 12 combinations of promoter and ORF into a low copy ARS/CEN plasmid. Wild-type *S. cerevisiae* strains were transformed with the individual plasmids carrying chimeric SUL constructs and competed against a plasmid-free strain to calculate the relative fitness coefficient of each strain in sulfate-limited media. Additionally, the non-chimeric alleles were also tested against a plasmid-free strain, with a total of 16 alleles tested.

The fitness coefficient values ranged from 0.2 to 38% after correcting for the cost of carrying a plasmid (-5.4% ± 0.59) as seen in **Fig 8**. When placed under the same promoter, the *SuSUL1* ORF now had a fitness advantage versus the *SuSUL2* ORF, opposite to the result obtained when each ORF was driven by its native promoter. All chimeras containing the promoter region of *SuSUL1* showed substantial decreases in fitness. This result suggests that expression differences between the two species may largely explain the differential fitness effects of the two *SUL1* genes. Interestingly, the chimeric allele containing the *SuSUL2* promoter with the *SuSUL1* ORF (*P_SuSUL1_-SuSUL1*) recapitulates the fitness effect of *ScSUL1*. Additionally, strains containing the promoter of *ScSUL1* or *ScSUL2* resulted in similar fitness patterns when paired with the three other ORFs, with the *ScSUL1* coding region yielding the highest relative fitness. However, when promoters of *SuSUL1* or *SuSUL2* were paired with the other three ORFs, we identified a different ranking of fitness patterns, with the *SuSUL1* coding region yielding the highest fitness. We did not attempt to further dissect these apparent epistasic interactions between the promoters and coding regions; however, such complex genetic interactions have been observed in other contexts [35–37].

**Fig 8.**
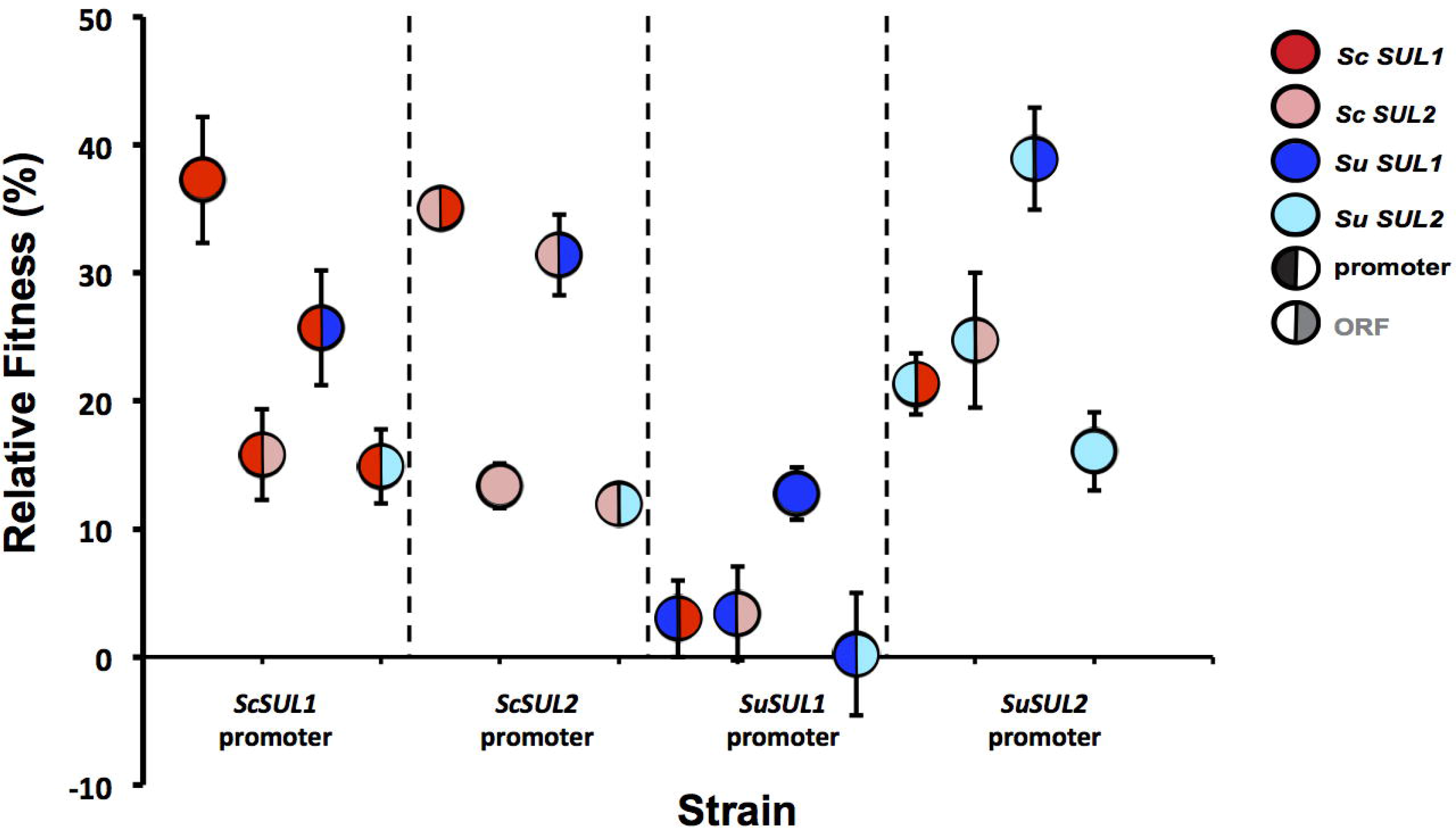
The promoter from *S. uvarum SUL1* reduces the fitness effect of *SUL1* and *SUL2* amplification. The relative fitness of 16 *S. cerevisiae* strains containing a plasmid including chimeric constructs of *SUL1* and *SUL2* alleles from *S. cerevisiae* or *S. uvarum*. The figure is split into quadrants by promoter; *ScSUL1, ScSUL2, SuSUL1, SuSUL2* (left to right) where each circle represents the chimeric construct. The left portion of the circle represents the promoter allele whereas the right portion represents the coding allele. The fitness was measured in the chemostat under sulfate-limited conditions against a GFP-marked lab strain. Each bar corresponds to the mean of 4 or more replicates ±SD.

Since the results from the chimeric constructs suggested that the promoter region is largely responsible for the differences in fitness, we sought to measure gene expression levels driven by each promoter. We used reverse transcriptase real time PCR (RT-PCR) to determine the expression level of *ScSUL1* under the control of all four promoters in *S. cerevisiae* strains grown at steady state in sulfate-limitation. We found that the expression level of the *ScSUL1* chimera with the promoter from *SUL1* from *S. uvarum (P_SuSUL1_-SCSUL1)* was significantly reduced in comparison to the other promoters (**Fig 9A**). We also found a modest correlation between expression level and the fitness value of each construct (R^2^=0.75) (**Fig 9B**). This result demonstrates that the differences between the fitness contributions of the two transporter genes may be due to gene expression differences.

**Fig 9.**
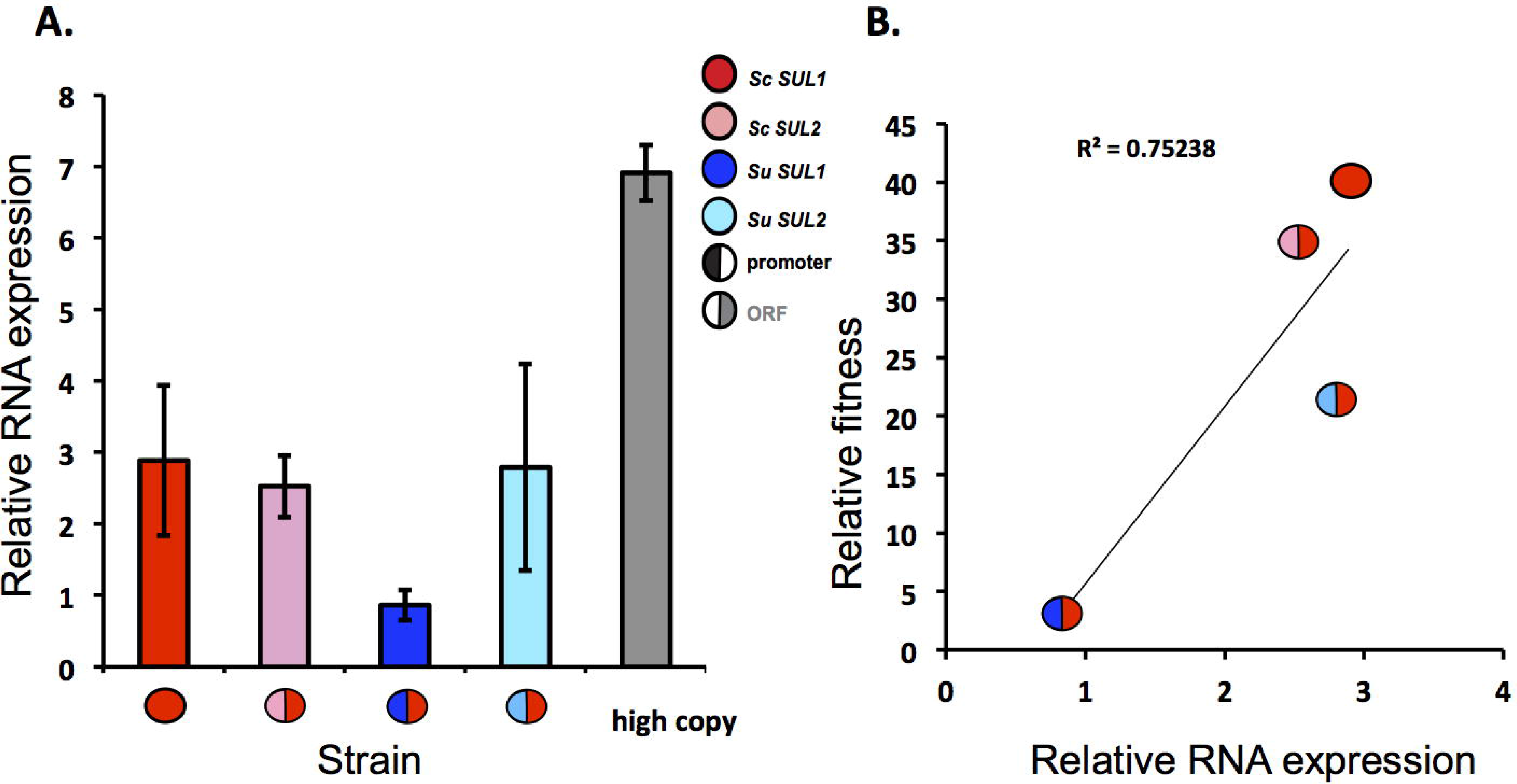
Reduced expression of chimeric construct with the *S. uvarum SUL1* promoter. A) The relative expression level of four *S. cerevisiae* strains containing a plasmid including the coding region of *S. cerevisiae SUL1* and the non coding region of *SUL1* or *SUL2* from *S. cerevisiae* or *S. uvarum*. Each split circle represents the chimeric construct. The left portion of the circle represents the promoter allele whereas the right portion represents the coding allele. The grey fitness value is a control strain of *S. cerevisiae* with five copies of *SUL1*. B) The relative expression of the construct correlates with the relative fitness of the strain (R^2^=0.7526)

## Discussion

In this work, we used comparative experimental evolution to investigate how paralog divergence influences adaptation to sulfate limitation across different species of yeast in the *Saccharomyces* clade. We identified differential sub-functionalization of gene duplicates between paralogs that encode sulfate transporter genes in *S. cerevisiae* and *S. uvarum*. Collectively, our results display an example of adaptation through two different evolutionary trajectories, likely driven by regulatory divergence.

We have shown *SUL1* amplification during long-term growth in sulfate-limited conditions occurs in all species tested in the *Saccharomyces* clade except *S. uvarum*. While the number of *S. paradoxus, S. mikatae*, and *S. uvarum* populations that were used for the laboratory evolution experiments was small (n=2-4), we have repeatedly identified *SUL1* locus amplifications in all evolution experiments of wild type *S. cerevisiae* (n=18/18). Therefore, it is surprising that even within two evolved populations of *S. uvarum*, we did not identify *SUL1* amplification, but instead identified *SUL2* locus amplifications in both populations after 500 generations. Additionally, two of the evolved populations in *S. paradoxus* and one population of *S. mikatae* amplified *SUL1*. The other populations that did not amplify *SUL1* or *SUL2* may contain other events that may be equally or more beneficial than either amplification, or may require additional time for the amplification event to occur and rise to high frequency (>200 generations). This point is further supported by additional evolution experiments we performed in *S. uvarum* for 200 generations where neither *SUL1* nor *SUL2* amplifications were detected, suggesting that amplification events are dynamic and may depend on longer time scales to occur and/or achieve high frequency. These findings also demonstrate that other means to achieve an increased fitness in sulfate limitation exist, since both *S. paradoxus* and *S. mikatae* are able to adapt to this condition without amplifying either of the SUL genes. Further work will be required to understand the genetic differences that mediate these other evolutionary trajectories.

Our results contribute to ongoing efforts to understand the mutations that drive adaptation, a long-standing question in evolutionary biology. There are examples of parallel molecular evolution that occur across genetic backgrounds for many traits [4,38–42], suggesting that genetic background plays a relatively unimportant role in determining the outcome of adaptation at the molecular level. A more recent study, however, tested how genetic differences between strains of bacteria influence their adaptation to a common selection pressure and found that parallel evolution was more common within-strains than between-strains, implying that genetic background has a detectable impact on adaptation [43]. Taken together, it is unclear to what degree genetic background impacts the mechanism and rate of adaptation to a novel selection pressure. Our study has identified differential locus parallelism between sulfate transporter loci in *S. cerevisiae* and *S. uvarum*, demonstrating one example where genomic background does substantially affect the outcome of evolution.

To further investigate the effect of genetic context and whether this was due to coding or non-coding variation, we generated chimeric alleles of promoter and coding regions between *S. cerevisiae* and *S. uvarum SUL1* and *SUL2* genes. We identified poor fitness outcomes associated with the non-coding region of the *SUL1* gene in *S. uvarum*, along with other complex interactions with the coding regions. These results suggest that the accumulation of mutations in the non-coding region of *S. uvarum SUL1* may have resulted in reduced expression, thus driving selection for *SUL2* amplification during adaptation of *S. uvarum* to sulfate limited conditions. Rich *et al* recently used a deep mutational scanning approach to identify the functional elements of the *ScSUL1* promoter that are crucial for growth in sulfate limitation [34]. This same approach could be applied to the promoter region of *SUL1* in *S. uvarum* to determine which sequences are responsible for these differences in activity.

Many studies have aimed to determine whether adaptation and phenotypic change typically occur from mutations in non-coding or coding regions in the genome [44–48]. In the case of gene duplicates, it has been proposed that their retention provides genetic redundancy, buffering the mutational space to either acquire new function, or to partition the ancestral function between duplicates. Gradual stochastic changes in expression level may lead to an eventual imbalance in the selective pressure between the two duplicates [49]. These gradual changes in gene expression may play a significant role in shaping the adaptive landscape over time, resulting in different adaptation outcomes across diverse genetic backgrounds. Our results provide an example of divergent adaptation through changes in expression of one duplicate in the *S. uvarum* lineage in the *Saccharomyces* clade. In the case of nutrient limitation, a simple modification in expression may be more likely to suffice, since the metabolic pathway for uptake and utilization already exists, and increasing uptake is a straightforward solution [4]. Alternatively, differential tradeoffs between toxic metal resistance and ion transport may exist between species and result in altered sulfur biosynthesis requirements to synthesize glutathione, a key factor in the cell's defense against oxidative stress and metal toxicity [50–52].

In addition to metal exposure, nutrient limitation is also a likely scenario experienced by wild and industrial yeast strains. Growing evidence suggests that domesticated *Saccharomyces* species have been exposed to sulfate related selective pressures through the selection for favorable characteristics associated with brewing. In lager brewing yeast, increased sulfite production is important for its antioxidant properties and for preserving favorable flavor profiles [53]. *Saccharomyces pastorianus* is a lager brewing species found only in the brewing environment and appears to be an allotetraploid hybrid between *S. cerevisiae* and *S. eubayanus* [54]. Interestingly, *S. pastorianus* carries inactive copies of *SUL1* from *S. cerevisiae* and *S. eubayanus*, while retaining functional copies of *SUL2* which have been shown previously to improve sulfite production when overexpressed [55–57]. Identifying the genetic basis of traits under selection in a particular environment may not only help highlight the emergence of new traits but also inform ways to engineer further improvement.

## Materials and Methods

### Yeast strains, plasmids and culture conditions

The strains used in this study are listed in **Table S1**. The *S. cerevisiae* strains used in this study were FY4 in the S288c background. The *S. uvarum* strains used were derived from the CBS1007 background. The *S. mikatae* strain was IFO 1815 and the *S. paradoxus* strain was CBS 432. The *N castellii* strain was CBS 4309. The *SUL1* and *SUL2* deletion strains were created in *S. cerevisiae* and *S. uvarum* by targeting 50bp upstream of the ATG and 100bp upstream of its stop codon. The deletions were confirmed with primers targeting approximately 175bp upstream of the ATG (**Table S3**).

To test the fitness due to the amplification of *SUL1* or *SUL2* from each species, we transformed DBY7283, a *ura3S. cerevisiaeMATα* strain, with a low-copy plasmid [58]. Phusion PCR was used to amplify 500bp upstream and 5bp downstream of the stop codon of *SUL1* and *SUL2* from *S. cerevisiae, S. uvarum, S. paradoxus*, and *S. mikatae*, and *SUL2* from *S. castelli*. Each *SUL1* and *SUL2* gene was blunt cloned into pIL37 using primers listed in **Table S3**. All plasmids used in this study are listed in **Table S2**. The haploid *S. cerevisiae* strain used in the competition experiments was a haploid FY MATa where the *HO* locus had been replaced with *eGFPas* previously described [26]. The diploid *S. cerevisiae* GFP+ strain was made by crossing the haploid FY MATα strain, where the *HO* locus had been replaced with *eGFP*, to a MATa FY strain. The *S. uvarum* GFP+ haploid strain was created by replacing the *HO* locus with eGFP by amplifying the NatMX-GFP construct from the plasmid YMD1139. The strain was verified using primers that target 600pb upstream of the *HO* locus. The fitness of the haploid *S. uvarum* GFP+ strain, YMD2869, was 0.388% +/− 0.33 (n=2). The fitness of the diploid *S. uvarum* GFP+ strain, YMD2869, was 2.33% +/− 0.19 (n=2). To directly compete two strains each containing an additional copy of either *SUL1* or *SUL2* from *S. cerevisiae* or *S. uvarum*, a GFP+ *ura3 S. cerevisiae* strain was transformed with plasmids containing either *SUL1* or *SUL2* from *S. cerevisiae* or *S. uvarum*. These GFP+ strains were used in a competitive assay against strains also containing additional copies of each gene.

The chimeric plasmids were created by amplifying 500bp upstream of the start codons of the *SUL1* and *SUL2* ORFs from *S. cerevisiae* and *S. uvarum* and cloning each upstream region into YMD2307 using primers with added SnaBI sites at the 3’ end (**Table S3**). Each plasmid was digested with SnaBI and *SUL1* or *SUL2* from *S. cerevisiae* or *S. uvarum* was ligated immediately adjacent to the previously cloned upstream region, creating a total of twelve different chimeric strains.

### Creation of hybrids

*de novo* hybrids between *S. uvarum* and *S. cerevisiae* were created by mating. Pulsed field gel analysis of the resulting strains confirmed the presence of both sets of chromosomes with no apparent size polymorphisms. Microarray analysis (see protocol below) of the hybrid DNA versus purebred DNA from each species also confirmed that these strains contained a complete haploid genome from each parent. Microarray data will be deposited in the Gene expression Omnibus (GEO) repository and in the Princeton Microarray Database.

### Microarray design

The *S. cerevisiae* and *S. uvarum* genomes were downloaded from the *Saccharomyces* Genome Database and concatenated to create a hybrid genome. The program Array Oligo Selector was used to design 70mers to each open reading frame in both genomes. Under the default stringency settings, 711 genes were too similar to another sequence in the combined genomes for a sufficiently unique oligo to be designed. For these cases, the program was rerun in the context of each single genome in order to provide more complete coverage of the purebred genomes. 485 genes were still too similar to other sequences in the single genomes to pass this test and were left off the array. The resulting 4840 *S. uvarum* and 6423 *S. cerevisiae* 70mers were purchased from Illumina.

### Microarray printing and preparation

70mer DNA was resuspended at 40 μM in 3X SSC and printed using a pin-style arraying robot onto aminosilane slides in a controlled-humidity environment. Slides were UV crosslinked at 70 mJ. On the day of hybridization, the slides were blocked by agitating for 35 minutes at 65C with 1% Roche blocking agent in 5X SSC and 0.1% SDS. Slides were then rinsed with water for 5 minutes and spun dry.

### Comparative genomic hybridization

Hybridization conditions were optimized to maximize specificity. DNA from *S. cerevisiae* was labeled with one fluor and DNA from *S. uvarum* labeled with another and competitively hybridized to the arrays under a variety of DNA quantity, hybridization volume and temperature, and wash stringency conditions. As expected because of the 2-tier design strategy, less than 5% (563/11263) showed evidence of cross-hybridization with signal significantly over background levels in both channels. These probes were filtered out of all hybrid datasets.

All microarray manipulations were performed in an ozone-free environment. 4 sμg DNA was sonicated to a size range near 1 kb then purified by Zymo DNA clean and concentrator columns. Labeling of 2 μg sonicated DNA was done by random-primed klenow incorporation of Cy-nucleotides either with the Invitrogen Bioprime kit according to the manufacturer's instructions, or with individually purchased reagents as previously reported [59]. The labeled reactions were purified by Zymo columns and measured for labeling yield and efficiency using a nanodrop spectrophotometer. 1 μg of each labeled DNA were mixed with Agilent blocking reagent and 2X hybridization buffer in a total volume of 400 μl, heated at 95C for 5 minutes, and hybridized to a prepared microarray using an Agilent gasket slide. Hybridizations were performed overnight at 65C in a rotating hybridization oven. Gaskets slides were removed in 1X SSC and 0.1 % SDS solution. Arrays were agitated for 10 minutes in a 65C bath of the same wash buffer, then washed on an orbital shaker for 10 minutes in a new rack in 1X SSC, ending with 5 minutes in 0.1 X SSC. Arrays were then spun dry and scanned in an Agilent scanner. The resulting images were analyzed using Axon Genepix software version 5. Complete microarray data will be available for download from the Princeton Microarray Database and GEO.

Data were linearly normalized and filtered for spots with intensity of at least 2 times over background in at least one channel. Manually flagged spots were also excluded. These filters were adequate to routinely filter out >95% of empty spots and retain >95% of hybridizing spots.

### Continuous culture evolution experiments

A single colony of *S. mikatae* and *S. paradoxus* and *S. uvarum* was inoculated into sulfate-limited chemostat medium with ura supplemented, grown overnight at 30°C, and 100 μL of the culture was inoculated into ministat chambers [25] containing 20 mL of the same medium at 30°C. After 30 hr, the flow of medium was turned on at a dilution rate of 0. 17 ± 0.01 hr^−1^. Four chemostats were inoculated from four independent clones for each species and cell samples (glycerol stock and dry pellet) were passively collected every day from fresh effluent for ~200 generations. DNA was isolated by a modified Smash-and-Grab protocol from each endpoint population [60]. Whole genome sequencing of the evolved and ancestral populations was performed as previously described

Longer term *S. uvarum* and hybrid evolution experiments were performed in ATR Sixfors fermentors modified to run as chemostats, as described [24], with the exception that *S. uvarum* populations were held at 25C. Prior experiments comparing this system with the ministat system demonstrated that they are nearly equivalent [25].

To determine if *SUL1* would amplify in *S. uvarum*, a single colony from a *sul2* strain was inoculated into sulfate-limited chemostat medium and inoculated from four independent clones into four ministat chambers as previously described. Array CGH was performed on the four populations after 260 generations using the ancestral strain as the reference (sul2Δ deletion strain).

Yeast samples for real-time PCR analysis were collected directly from the culture vessels, when the cultures reached steady state (approximately 3 days at ~25 generations). The cells were filtered on Nylon membrane (0.45μm pore size) and immediately frozen in liquid nitrogen and stored at −80°C until RNA extraction.

### Competition experiment

The pairwise competition experiments were performed in ministats. Each competitor strain was cultured individually. Upon achieving steady state, the competitors were mixed in the indicated ratio. Each competition was conducted in two biological replicates for 15 generations after mixing. Samples were collected and analyzed three times daily. The proportion of GFP+ cells in the population was detected using a BD Accuri C6 flow cytometer (BD Biosciences). The data were plotted as ln[(dark cells/GFP+ cells)] vs. generations. The relative fitness coefficient was determined from the slope of the linear region by the use of linear regression analysis.

### Total RNA extraction and Quantitative RT-PCR

RNA was extracted from the filtered sample by acid phenol extraction and quantified using the nanodrop. 90μg of RNA was cleaned-up using the Qiagen RNA easy kit according to the manufacturer's instructions (Qiagen). Contaminating DNA was removed by using Rapid DNase out removal kit on 2μg of RNA in a 100μL reaction (Thermo).

Oligonucleotides for real-time PCR are listed in **Table S3**. One microgram of total RNA was reverse-transcribed into cDNA in a 20 μL reaction mixture using the SuperScript VILO cDNA synthesis kit (Life). The cDNA concentrations were then analyzed using the nanodrop. For the RT-PCR, each sample was tested in triplicate in a 96-well plate using SYBGR. The reaction mix (19 μL final volume) consisted of 10 μL of LightCycler 480 SYBR Green I Master (Roche), 2 μL of each primer (5 mM final concentration), 5 μL of H_2_O, and 1 μL of a 1/100 dilution of the cDNA preparation. The absence of genomic DNA in RNA samples was verified by real-time PCR using the DNase free RNA. A blank was also incorporated in each assay. The thermocycling program consisted of one hold at 95°C for 4 min, followed by 50 cycles of 10 sec at 95°C and 45 sec at 56°C. The quantification of the expression level of *SUL1* was normalized with *ACT1*.

### Nextera libraries and whole-genome sequencing

Genomic DNA libraries were prepared for Illumina sequencing using the Nextera sample preparation kit (Illumina). Barcoded libraries were quantified on an Invitrogen Qubit Fluorometer and submitted for 150 bp paired end sequencing on an Illumina HiSeq 2000. Read data have been deposited at the NCBI under BioSample accessions: SAMN04211938, SAMN04211939, SAMN04211940, SAMN04211941, SAMN04211942, SAMN04211943, SAMN04211944, SAMN04211945, SAMN04211946, SAMN04211947, SAMN04211948, SAMN04211949 and SAMN04211950. The reads were aligned against the reference strain of *S. uvarum* (CBS 7001), *S. mikatae* (IFO 1815), and *S. paradoxus* (CBS 432) using Burrows-Wheeler Aligner [61]. The sequence coverage of the nuclear genome ranged from 70 to 300x. Copy-number variations (CNVs) were detected by averaging the per-nucleotide read depth data across 100bp windows. For each window, the log_2_ratio in read depth between the evolved and parental strain was calculated. The copy number was calculated from the log_2_ratios and plotted using the R package DNA copy [62].

## Acknowledgements

We thank members of the Dunham lab for helpful discussions and Celia Payen and Caiti Smukowski Heil for their helpful comments on the manuscript. We also thank Lory Henderson for her help characterizing the growth of the *S. uvarum* SUL deletion strains, as well as Noah Hanson with his help with the competition assays. Additionally, we thank Rachel Youngblood for her assistance with making sequencing libraries of the evolved *S. mikatae* and *S. paradoxus* strains. Thanks to Yixian Zheng and Doug Koshland for contributing to the initial experimental design, creating yeast strains, and purchasing the oligonucleotides used for the microarrays. This work was funded by NSF grants 1516330, 1243710, and 1120425, and NIH grant R01 GM094306. MD is a Senior Fellow in the Genetic Networks program at the Canadian Institute for Advanced Research and a Rita Allen Foundation Scholar. MS was funded by NSF GSRF DGE-1256082 and the Robert D. Watkins Graduate Research Fellowship from the American Society for Microbiology. ABS was funded by F30CA165440 and IL by F32GM090561. Funding was also from P50 GM071508 to the Lewis-Sigler Institute and from the Howard Hughes Medical Institute to Doug Koshland and Yixian Zheng.

## Supplemental Legends

**S1 Table. List of strains**

**S2 Table. List of plasmids**

**S3 Table. List of primers**

**S1 Fig. Chromosome X copy number plots of two evolved populations of *S. uvarum*.**

**S2 Fig. Chromosome XII copy number plots of four evolved *sul1*Δ *S. cerevisiae* populations**

**S3 Fig. Chromosome II copy number plots of 16 evolved hybrid clones**

**S4 Fig. Chromosome XII copy number plots of 16 evolved hybrid clones**

**S5 Fig. Whole genome copy number plots of four evolved *S. paradoxus* populations**

**S6 Fig. Whole genome copy number plots of four evolved *S. mikatae* populationsCopy Number**

